# Digestive exophagy of Bacterial Biofilms by an Amoeba Predator is Mediated by Specific Biofilm Recognition

**DOI:** 10.1101/2022.09.24.509356

**Authors:** Eva Zanditenas, Meirav Trebicz-Geffen, Laura Domínguez-García, Diego Romero, Ilana Kolodkin-Gal, Serge Ankri

**Affiliations:** Department of Molecular Microbiology, Ruth and Bruce Rappaport Faculty of Medicine, Technion, Haifa, Israel; Departamento de Microbiología, Instituto de Hortofruticultura Subtropical y Mediterránea ‘La Mayora’, Universidad de Málaga-Consejo Superior de Investigaciones Científicas (IHSM-UMA-CSIC), Universidad de Málaga, Málaga, Spain; Department of Plant Pathology and Microbiology, the Robert H. Smith Faculty of Agriculture, Food & Environment, The Hebrew University of Jerusalem, Rehovot, Israel

## Abstract

The human protozoan parasite *Entamoeba histolytica* is responsible for amebiasis, a disease endemic to developing countries. *E. histolytica* trophozoites are released from the cysts to colonize the large intestine, where they primarily feed on bacterial cells. In these scenarios, bacterial cells form aggregates or structured communities too large for phagocytosis. Our results show that *E. histolytica* can degrade pre-established biofilms of *Bacillus subtilis* and *Escherichia coli* in a dose- and time-dependent manner. Surprisingly, trophozoites incubated with *B. subtilis* biofilm exhibit a unique transcriptome signature compared to those incubated with planktonic cells or without bacteria. Biofilm-induced genes include cysteine proteases (CPs), and the general inhibition of CPs by E64D or by the use of specific small-RNA (sRNA)-based RNA interference impairs the degradation of biofilms by *E. histolytica*. The degradation of *B. subtilis* extracellular matrix (ECM) protein TasA by CPs is associated with partial biofilm digestion and activation of the stress response in the interacting *B. subtilis* cells. The interaction with *B. subtilis* biofilms was also associated with lower levels of oxidoreductases. Oxidoreductase downregulation can be a readout of the embedding of *E. histolytica* trophozoites within the biofilm-produced extracellular matrix, reducing their exposure to oxidative stress (OS). Our results indicate that parasites may digest biofilms by a controlled mechanism of digestive exophagy as secretion of digestive enzymes as a conserved mechanism for biofilm degradation allows phagocytic digestion of biofilm cells. Furthermore, the partially digested biofilms can serve as an unexpected shield protecting parasites from oxidative environments and thereby may regulate the persistence and virulence of the parasite.

## Main Text

*Entamoeba histolytica* is a protozoan parasite responsible for amebiasis, a wildly prevalent human intestinal disease in developing countries. Transmission of amebiasis occurs mainly through contaminated food or water with feces containing *E. histolytica* cysts, which is one of two forms of the parasite^1^. On entering the host intestine, the cyst, the resistant form of the parasite, undergoes a process called excystation, during which trophozoites, the parasite’s vegetative form, are released. In most of the cases of infection, these trophozoites feed on bacterial microbiota or cellular debris in the large intestine without causing symptoms. In symptomatic infection characterized by bloody diarrhea, the parasite damages the mucus layer and intestinal epithelial cells, causing an inflammatory response that involves the recruitment of neutrophils and macrophages and the release of reactive oxygen species and nitric oxide by these immune cells^2^.

An estimated 10^14^ microorganisms inhabit the large intestine. There has been increasing recognition that gut bacteria play a key role in the development of amebiasis. (for a recent review see^3^). For instance, cultivating *E. histolytica* with *E. coli* O55 can increase *E. histolytica’s* virulence, which is dependent on the contact between the amoeba and bacteria^4^. Furthermore, *E. histolytica* trophozoites showed greater resistance to oxidative stress (OS) after incubation with *E. coli* O55 ^5^. Infection with *E. histolytica* can cause dysbiosis marked by reduction of Lactobacillus and Bacteroides and increase in Bifidobacterium ^6^. These studies have examined the interaction of planktonic bacteria with *E. histolytica* trophozoites; however, the interaction of this parasite with bacterial biofilms remained poorly characterized. This uncharted area is of high ecological and clinical relevance as in the intestinal tract, and bacteria reside as complex microbial communities that are not planktonic. Instead, microbiota members form higher order structures frequently termed biofilms, as they are embedded in complex, self-produced polymeric matrices, adherent to each other and surfaces or interfaces, and have enhanced antimicrobial resistance, virulence, and quorum sensing capacities ^7,8^.

*Bacillus subtilis* is a genetically manipulatable model of a biofilm-forming bacterium, from the *firmicutes* phylum. This Gram-positive bacterium that lives in soil, is a proficient biofilm and the molecular mechanisms responsible for biofilm formation by this bacterium while grown in isolation are well established ^9^. *B. subtilis* is found in the human gastrointestinal tract, as it is currently widely used as a probiotic ^10^ intended to promote digestive health and a healthy immune system in both adults and children ^11^. *B. subtilis* forms robust biofilms as structured floating pellicles growing on the surface of liquid cultures and architectonically complex colonies on agar plates ^12^. *B*. subtilis biofilm formation is triggered by external factors like low oxygen or nutriment depletion and is governed by the master regulator Spo0A^13^. An initial step in biofilm formation is the transition of the planktonic cells to sessile state triggered by the downregulation of flagella genes expression ^14^ and the upregulation of genes responsible for producing extracellular matrix (ECM). The most extensively abundant components of biofilm organic extracellular matrix are carbohydrate-rich polymers (i.e., extracellular polysaccharides or exopolysaccharides), and proteins ^7^. TasA and TapA are the proteinous components of the extracellular matrix and are essential for the biofilm’s rigidity, and complex 3D architecture ^7^. TasA forms amyloid fibers ^15–17^ that are attached to the cell wall and, in conjunction with other extracellular components, promote cell-cell adhesion ^16,18^. BslA is secreted at the end of the biofilm formation process in order to form a hydrophobic layer that confers broad-spectrum resistance to antimicrobials ^19^. While *B. subtilis* matrix was shown to protect cells from antibiotics and abiotic stressors^9,19^, it remained unknown whether this matrix, as well as matrices produced by other bacteria can protect biofilm cells from predation and phagocytosis.

The most well characterized resident of the GI is *E. coli*. Foodborne outbreaks associated with fresh produce have been increasing, with *E. coli* being the most common pathogen associated with them^20^. A notorious example is the major outbreak of the hemolytic-uremic syndrome caused by horizontally transferred Shiga-toxin-producing *E. coli O104:H4* in Germany and France that was associated with sprout consumption^21^. *E. coli* is a Gram-negative bacterium residing in the mucosal layer of mammalian gastrointestinal tract where it is the most abundant facultative anaerobe. The biofilms of *E. coli* depend on a functional amyloid curli ^22^ that enables cell-cell adherence, adherence to vegetables^23^ and to a mammalian host^24^. Two divergent operons *csgBAC* (<u>c</u>urli-<u>s</u>pecific <u>g</u>ene) and *csgDEFG*, are necessary and sufficient to produce curli ^25^. In *E. coli*, curli fibers compose up to 85% of the biofilm biomass ^26^, spatially expressed in the wrinkles of structured colonies ^27^ and often form an interwoven mesh that cradle the individual bacterial cell ^28^. The curli fibers from *Enterobacteriaceae* are the foremost studied functional amyloids ^22^. In both *E. coli* and *B. subtilis*, biofilm biology critically depends on amyloid formation. Here, we aimed to characterize the molecular mechanisms of interaction between parasites and biofilms sharing the same habitat, using the genetically manipulatable *E. histolytica* and proficient biofilm formers *B. subtilis* and *E. coli*. These bacterial represent a Gram-positive probiotic strain ^29^, and a Gram-negative enteric pathogen^30^, respectively. Once bacteria form biofilms, simple phagocytosis is no longer feasible as biofilms are objects too large for phagocytosis. Yet, the interactions of small protozoan predators and biofilms were poorly studied^31,32^. This study describes the mechanisms that promote a predator-prey interaction between assembled biofilms and their predator. Specifically, we uncovered that the transcriptome architecture of *E. histolytica* reflects a specific recognition/response of/to *B. subtilis* biofilm, which differs from planktonic *B. subtilis* cells, the role of CPs in the degradation of the biofilm, and the protective role of the *B. subtilis* biofilm against OS. These unexpected outcomes that highlight the importance of studying the *amoeba*-bacteria interactions while considering biofilms as the target prey.

## Results

### 1- *E. histolytica* trophozoites degrade *B. subtilis* biofilms in a dose and time dependent manner

Using biofilms where the matrix protein TasA was fused to mCherry (to specifically label biofilm cells and their assembled matrix)^33^ and GFP labeled amoeba we explored the nature of the interaction between the partners. When *E. histolytica* interacts with *B. subtilis* biofilms, the parasites first adhere to the surface of the biofilms (Fig. 1A). With time, the penetration of parasites into the biofilm is observed (Figs 1B and S7)). We therefore asked whether *E. histolytica* trophozoites are capable to degrade the assembled biofilms to promote their invasion/penetration.

**Figure 1:**
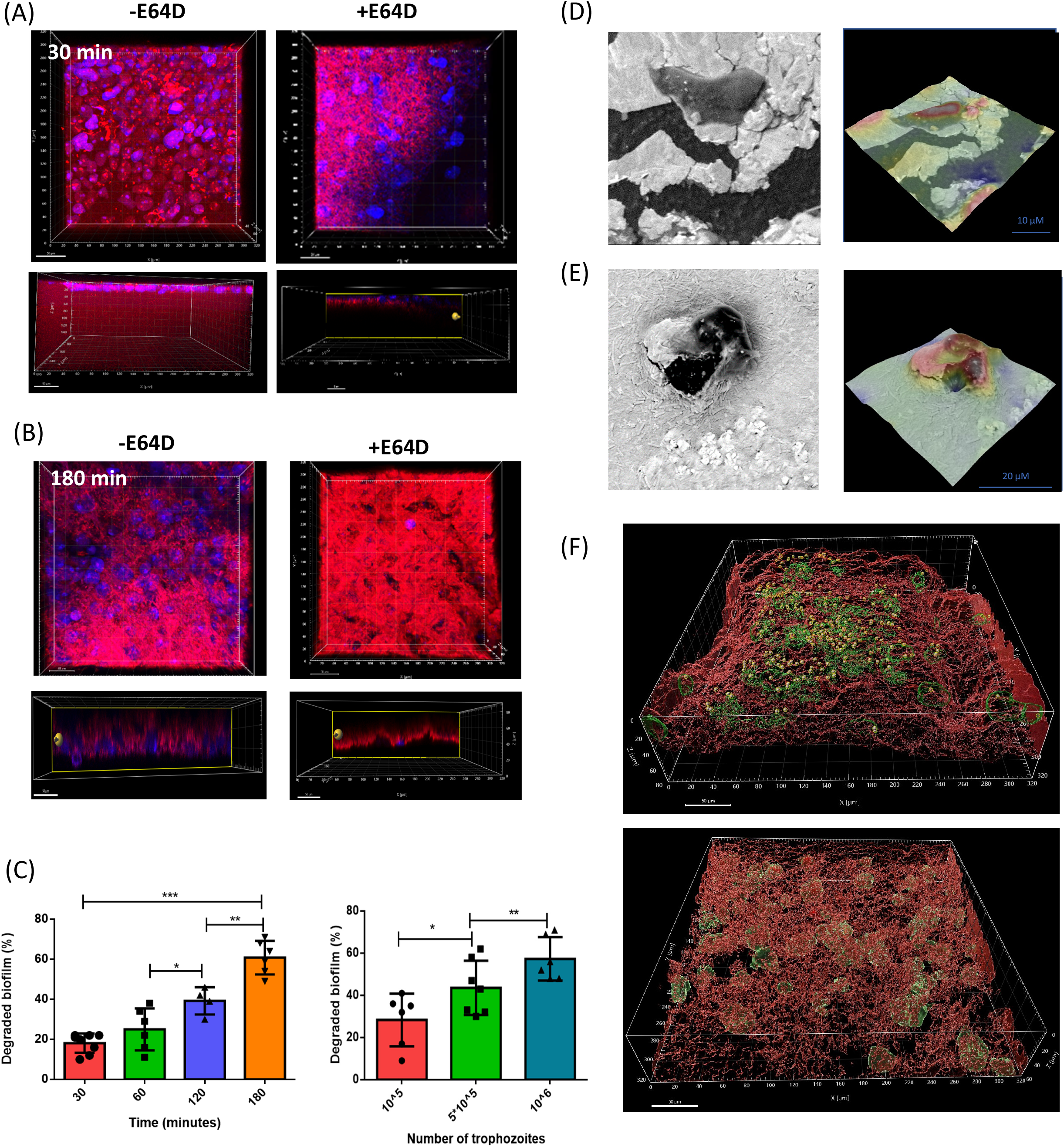
The destruction of *B. subtilis* biofilms by *E. histolytica*. (A) Confocal microscopy (X30) of *B. subtilis* biofilm that expresses TasA-mCherry after 30 minutes of incubation at 37°C with *E. histolytica* trophozoites (stained with DAPI) and treated with or without E64D. Upper panel shows top view and lower panel shows side view. (B) Images were obtained as in A for 180 minutes incubation (C) *B. subtilis* biofilm destruction by *E. histolytica* trophozoites in a time and dose dependent manner. student test, * p-value less than 0.05 and ** p-value less than 0.01. Data represent average of results from 3 biological replicates (D) Electron microscopy images of trophozoites present on the biofilm. Upper left panel: Trophozoites (in black) are present on the surface of the biofilm (in grey). (E) Upper right panel: heat map of trophozoites (in red) on the biofilm (in green) demonstrates degraded areas. Down left panel: trophozoites (in black) entering the biofilm embed within the biofilm. Lower right panel: destruction of the biofilm located below the parasite. (F) 3D model of trophozoites (stained with DAPI) treated (top panel) or not with E64D (lower panel) in the biofilm, that expresses TasA-mCherry (using Imaris software).

We evaluated the degradation of *B. subtilis* biofilms constitutively expressing green fluorescence protein (GFP) by measuring a decrease in fluorescence of the biofilm upon interaction with the parasite. The biofilm was degraded by 5.10^5^ trophozoites in a time-dependent manner, with maximum degradation occurring after three hours of incubation (Fig. S1). Moreover, we investigated the effect of trophozoites number on degradation of biofilms with trophozoites. Following 3 hours of incubation at 37°C, 10^5^ and 5.10^5^ trophozoites degraded 28% and 47% of the biofilm, respectively. In the presence of more trophozoites (10^6^) to the biofilm, the degradation rate increased to 57%, suggesting that the system is saturated (Fig. 1C).

To further evaluate whether the interaction of trophozoites with biofilms is capable of destroying the biofilms, we used scanning electron microscopy to identify the parasites with certainty (Fig. 1D). Consistent with our destruction assays we could detect parasite that irreversibly attach to the biofilm surface (Fig. 1D) but also parasites generating a clear noticeable crack beneath them read by the 3D reconstruction as an empty area (Fig 1E). Indeed, trophozoites inducing the local degradation of the ECM were frequently hidden beneath biofilm surface (Fig. 1F). Furthermore, the trophozoites were capable of predating the biofilm cells as these TasA expression cells were imaged within the trophozoites throughout the film (Fig. 1F). 3D surface reconstruction confirmed the dynamic of the invasion of the trophozoites into the biofilm (data not shown).

An overall examination of the biofilms revealed a clear loss of structural topography upon interaction with the parasite (Fig. 1A and Fig. S2). Consistent with our observation that *B. subtilis*, serves as a prey for the trophozoites, we tested whether cells from the partially degraded biofilms experience an elevated stress following their exposure to the predator. Indeed, the extracts from trophozoites were sufficient to induce both the general stress response and the cell wall stress response in biofilm cells (Fig. 2B, C).

**Figure 2:**
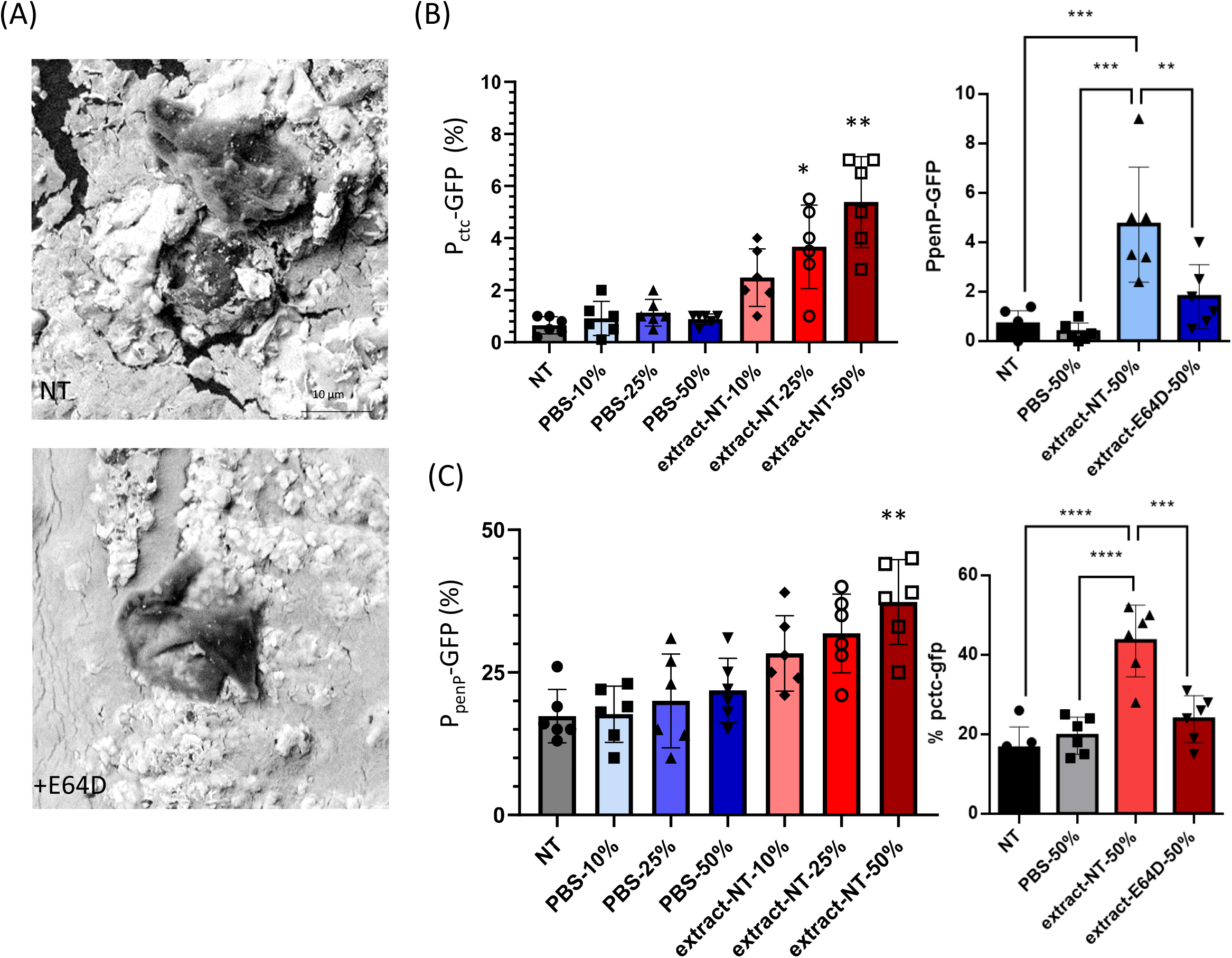
*B. subtilis* biofilm cells sense predation by *E. histolytica*. (A) Scanning electron microscopy of the overall biofilm topography in tropozoites either untreated or treated with E64D (B) A WT *B. subtilis* strain harboring Pctc-GFP (General stress response) or PpenP-gfp (SigW mediated cell wall stress) were analyzed in the presence and absence of increasing concentrations of *E.histolytica’s* extract using flow cytometry. Colonies were grown on B4 medium and incubated at 30 °C. Data were collected from 24 h post inoculation in MSgg, 100,000 cells were counted. Y-post-inoculation) the % of cells expressing the reporters, graphs represent mean ± SD from three independent experiments (n = 6). **<0.01, ***<0.001 and ****<0.0001 (C) Same as C, but extracts were collected from with untreated amoebae or amoeba’s treated with E64D. **<0.01, ***<0.001 and ****<0.0001

### 2. The transcriptome architecture reflects differential recognition of biofilms and planktonic cells

We used RNA sequencing (RNA-seq) to examine the mechanism of this antagonistic interaction and subsequent degradation of *B. subtilis* biofilms by *E. histolytica* trophozoites. Transcriptomics of wild type trophozoites (WT), WT incubated with planktonic *B. subtilis* (pB) and WT incubated with *B. subtilis* biofilm (bB) was compared.

In the WT+pB versus WT comparison 157 and 199 transcripts were respectively induced or repressed (table 2). In the WT+bB versus WT comparison 515 versus 543 transcripts were respectively induced or repressed (table 2). In the WT+ bB versus WT+ pB comparison 241 versus 168 transcripts were respectively induced or repressed (table 2).

A hierarchical clustering analysis presented as a heatmap indicates that WT and WT+pB conditions cluster together whereas the WT+pB condition forms an independent cluster (Fig. 3A). Volcano plots indicated a minor/modest change of the transcriptome architecture of trophozoites interacting with planktonic *Bacillus subtilis cells*. In contrast, the differences between biofilm interactors and non-interacting trophozoites (control) were larger and included multiple genes which were both significantly changed (Fig. 3B and S3). Similar trends were observed when trophozoites that interact with planktonic cells were compared against trophozoites that interact with assembled biofilms (Fig. S3). These results indicate that *E. histolytica* cells respond differently to the same bacterium either grown as planktonic or as a biofilm, and that biofilms have unique distinguishable features recognized by the parasites.

**Figure 3:**
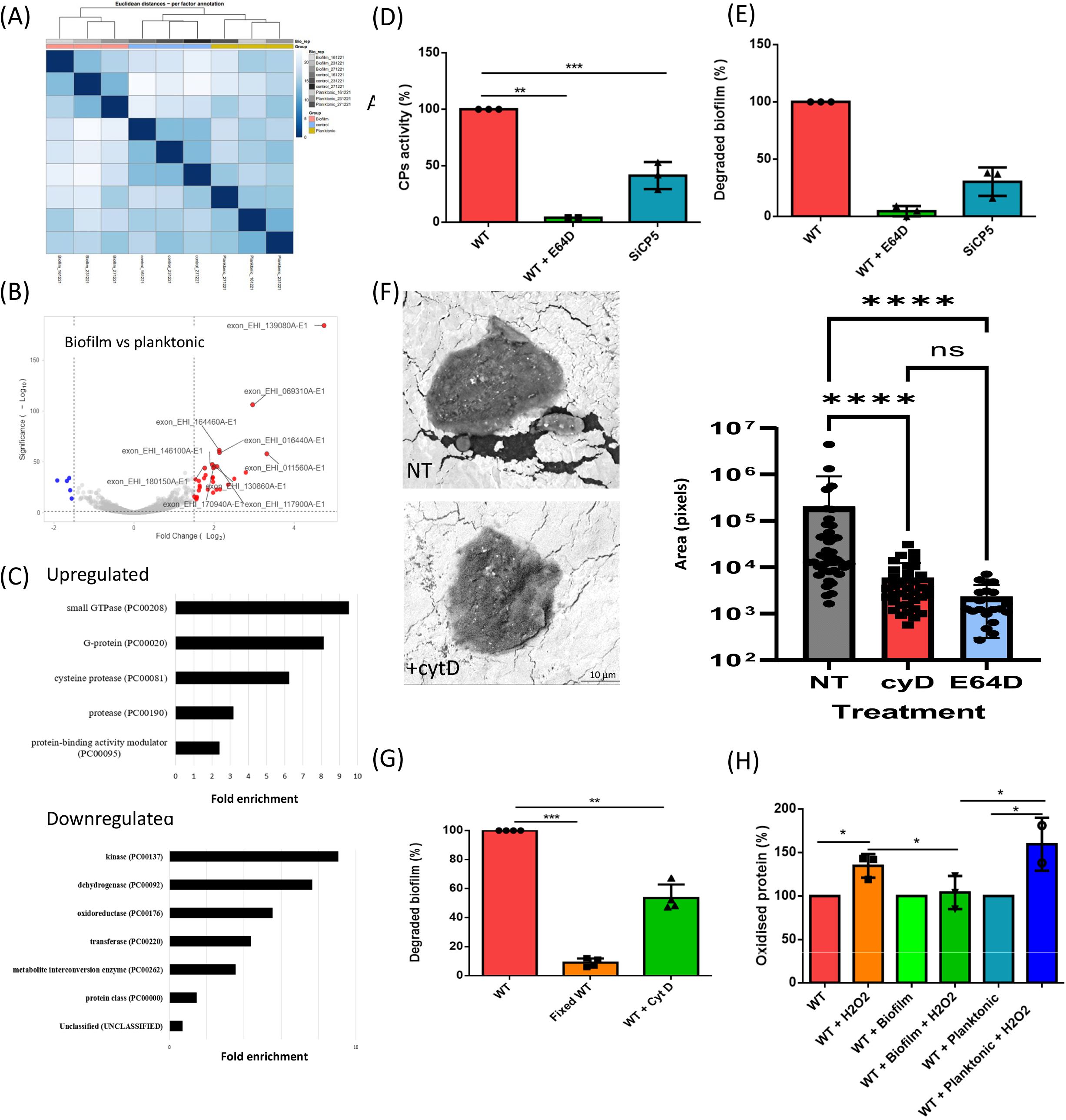
Specific recognition of *B. subtilis* biofilm cells by *E. histolytica*. (A) Heat-map of the transcriptomic results showing the Euclidean distance between control trophozoites and trophozoites incubated with *B. subtilis* planktonic or biofilm form. Darker color indicates stronger correlation. (B) Volcano plot of the trophozoites incubated with *B. subtilis* planktonic form VS trophozoites incubated with *B. subtilis* biofilm (C) PANTHER statistical overrepresentation test of upregulated and downregulated genes in AFAT. (D) CPs activity in control trophozoites (WT), E64D treated trophozoites and in siCP5 trophozoites. CP activity was measured with as substrate. (Student test, p-value <0.001). Data represent averages of results from 3 biological replicates. (E) *B. subtilis* biofilm degradation by control trophozoites (WT), E64D treated trophozoites and silenced CP5 trophozoites (siCP5) after 2 hours of incubation at 37°C. (Student test, ** p-value less than 0.01. *** p-value less than 0.001). Data represent averages of results from three biological replicates. (F) Scanning Electron Microscopy of a CytD or E64D treated trophozoites (left panel) and quantification of the cracked areas (right panel) in the indicated backgrounds from scanning electron Microscopy (n=5 fields). Crack size was analyzed by imageJ software. (G) *B. subtilis* biofilm degradation by control (live) or paraformaldehyde fixed trophozoites and by Cyt D treated trophozoites. (** p-value less than 0.01. *** p-value less than 0.001.). Data represent averages of results from three biological replicates (H) Quantification of oxidized proteins in trophozoites incubated with planktonic or biofilm *B. subtilis* and exposed to H_2_O_2_ (2.5 mM, 30 min). The data have been normalized using total protein normalization. Student test, ** p-value less than 0.01. Data represent averages of results from two biological replicates.

To test how *E. histolytica* can distinguish *B. subtilis* biofilm cells from planktonic cells, the differentially regulated genes in WT+ bB versus WT+ pB were classified, according to the protein class they encode, using PANTHER. The categories for functional classification of genes upregulated in WT+ bB versus WT+ pB are shown in Fig 2C. The most abundant classes are metabolite interconversion enzyme (PC00262) such as AIG1 family protein (table 3), protein-modifying enzyme (PC00260) such as cysteine proteinase A-4 (EHI_050570) and protein-binding activity modulator (PC00095) such as Rho family GTPase (EHI_180430). Of the upregulated genes in WT+ bB versus WT+ pB, genes that encode for small GTPase (PC00208) such as AIG1 family protein (EHI_136940) and cysteine protease (PC00081) such as cysteine proteinase (EHI_151440), are significantly enriched according to the PANTHER statistical overrepresentation test.

The categories for functional classification of genes downregulated in WT+ bB versus WT+ pB are shown in Fig. 2C. The most abundant class of gene encoded proteins are protein modifying enzyme (PC00260) such as thioredoxin reductase (EHI_155440) and metabolite interconversion enzyme (PC00262) such as malate dehydrogenase (EHI_092450). Of the downregulated genes in WT+ bB versus WT+ pB, genes that encode for kinase (PC00137) such as Pyruvate, phosphate dikinase (EHI_009530) and dehydrogenase (PC00092) such as NAD(FAD)-dependent dehydrogenase (EHI_099700) are significantly enriched according to the PANTHER statistical overrepresentation test.

The analysis of additional comparisons (WT+ pB versus WT and WT+ bB versus WT) is presented in Figs S4 and S5.

### 3. A role for cysteine proteases and cytoskeleton remodeling in the predation of *B. subtilis* biofilm cells by *E. histolytica*

A close examination of the proteinous matrix that surrounds *E. histolytica* (Figs 1B and S2) reveled areas of clearance of these proteinous components. Indeed, many cysteine protease genes are upregulated in *E. histolytica* trophozoites exposed to *B. subtilis* biofilms (table 2 and Fig 3C), suggesting that these enzymes are involved in the degradation of the biofilm. In order to test this hypothesis, *E. histolytica* trophozoites were treated overnight with the cell permeable cysteine protease inhibitor, E64D (10 μM). Incubation of trophozoites with E64D for 24 hours strongly inhibited cysteine protease activity (Fig. 3D) but has only a modest effect on the parasite (around 70% of the trophozoites are still viable). The ability of E64D-treated trophozoites to penetrate inside the biofilm, partially degrade biofilms of *B. subtilis* and phagocyte bacteria detached from the biofilm was also severely impaired (Fig 1A,B,F, S6,7 and 3E). EhCP5 (EHI_168240) is a main amebic virulent factor present on the surface of the parasite and also secreted ^34^. Even though EhCP5 expression was not increased in trophozoites exposed to *B. subtilis* biofilms, the presence of EhCP5 at the parasite’s surface suggests that this protease may play an important role in the early stage of biofilm degradation. In order to test this hypothesis, we used RNA interference gene silencing ^35^ to downregulate EhCP5 expression. The downregulation of EhCP5 expression in the EhCP5–silenced strain was confirmed by Q-RT-PCR (Fig. S8). When compared to the wild type parasite, EhCP5-silenced trophozoites show significantly decreased CP activity and ability to degrade *B. subtilis* biofilm (Fig 3D and E). These results strongly suggest that EhCP5 is involved in the degradation of *B. subtilis* biofilm. In agreement with the antagonistic effects of CPs on *B. subtilis* biofilm cells, the fold induction of stress genes in the presence of CP inhibitor was reduced (Fig. 2D and E).

Actin (EHI_164440; EHI_164430) and EF-hand calcium-binding domain containing protein (EHI_016120) among genes upregulated in *E. histolytica* trophozoites exposed to *B. subtilis* biofilms (table 2), suggests that cytoskeleton remodeling is involved in the degradation of the biofilm by the parasite. The hypothesis was tested by exposing trophozoites to cytochalasin D (Cyt D) (5 μM), an inhibitor of actin polymerization ^36^, before incubation with the biofilm. Trophozoites incubated with Cyt D failed to induce substantial cracks in the biofilm matrix and had their ability to penetrate inside the biofilm impaired and they degraded the biofilm two times less than those incubated without it (Figure 3F and G). Consistently, the biofilm degradation was more modest and manifested with less constrained degraded areas in trophozoites treated with CytD compared with CPs (Fig 3F). Based on these results, cytoskeleton remodeling also contributes to the entry inside the biofilm and its degradation.

### 5. The invasion into *B. subtilis* biofilms protects *E. histolytica* against Oxidative Stress

Our previous work support a role of bacterial metabolites in protecting *E. histolytica* against to reactive oxygen species (ROS) (for a recent review see ^3^). We observed that *E. histolytica* trophozoites penetrate biofilms and become embedded within them (Figs 1ABF). These findings raise a hypothesis that the parasites use the biofilm as a protective layer, potentially to ROS. Several genes involved in *E. histolytica* response to OS like thioredoxin reductase (EHI_155440) ^37^, EhNO1 (EHI_110520) (EhNO1 is mainly involved in ferric reduction) ^38^ or the DNA damage recognition like the excision repair protein RAD23 (EHI_001400) ^39^ have their level of expression downregulated in *E. histolytica* trophozoites exposed to *B. subtilis* biofilm (table 2). These data suggest that the biofilm protects the parasite against OS. In order to test this hypothesis, trophozoites alone or trophozoites in presence of planktonic or biofilm form of *B. subtilis* were exposed to H_2_O_2_-induced OS (Fig 3H and Fig. S9). The level of oxidized proteins (OXs) in the parasite was determined by immunoblot detection of carbonyl groups introduced into proteins by OS. In the presence of H_2_O_2_, OXs in control trophozoites were significantly higher than OXs in trophozoites incubated with either the planktonic or biofilm form of *B. subtilis* prior to their exposure to H_2_O_2_. A significant reduction in the level of OXs was observed in trophozoites in *B. subtilis* biofilms compared to trophozoites present outside the biofilms (Fig 3H). These results are consistent with the low oxygen concentrations measured within microbial biofilms^40^.

### 5. The role of the bacterial matrix protein TasA as a substrate for the binding of and degradation by *E. histolytica* trophozoites

In *E. histolytica*, the Gal/GalNAc lectin, composed of both heavy and light subunits, plays an important role in binding to mucins, host cell receptors, and bacteria^41^. As we could detect adhesion of the parasites to the biofilms (Fig. 1ADE), we explored the component that mediates the binding of the parasite to the biofilm. The presence of a gene encoding the Gal/GalNAc lectin heavy subunit (EHI_046650) among genes that are upregulated in *E. histolytica* trophozoites exposed to *B. subtilis* biofilm (table 2) suggests that the Gal/GalNAc lectin is involved in the binding of the parasite to the biofilm. To test this hypothesis, *E. histolytica* trophozoites were exposed to the biofilm in presence or absence of galactose (2%) or asialofetuin (0.05%) which act as competitive inhibitors to reduce the parasite’s Gal/GalNAc-lectin-dependent attachment to cells^42^. We found no significant differences in the degradation of *B. subtilis* biofilm by trophozoites incubated with and without galactose (Fig. 4A). These results indicate that the Gal/GalNac receptor on the surface of the parasite is not involved in binding to the *B. subtilis* biofilm and degrading it. Some bacteria, such as *Escherichia coli* and *Serratia marcescens* strains with mannose-binding components on their cell surfaces, bind to mannose receptors on amoebic membranes ^43^. By using mannose as a competitor, we tested if *E. histolytica* binds to *B. subtilis* biofilms via mannose. There was no significant difference in the degradation of *B. subtilis* biofilm by trophozoites incubated with or without mannose (2%) (Fig. 4A). These results indicate that mannose receptors on the parasite’s surface are not involved in binding to the *B. subtilis* biofilm and degrading it. In order to gain further insight into the mechanism of *E. histolytica* binding to *B. subtilis* biofilms, we used the planktonic form of *B. subtilis* as competitor. The planktonic form of *B. subtilis* does not impair the degradation of *B. subtilis* biofilms by *E. histolytica* (Fig. 4A). These results suggest that the parasite is binding to a specific component of the biofilm. To test whether TasA is involved in the binding of the parasite to the biofilm, we used a pure TasA as a competitor. We observed that TasA (10 μg) significantly reduced the ability of the parasite to attach to the biofilm (Fig 4A). This result strongly suggests that the parasite binds to the biofilm *via* TasA.

**Figure 4:**
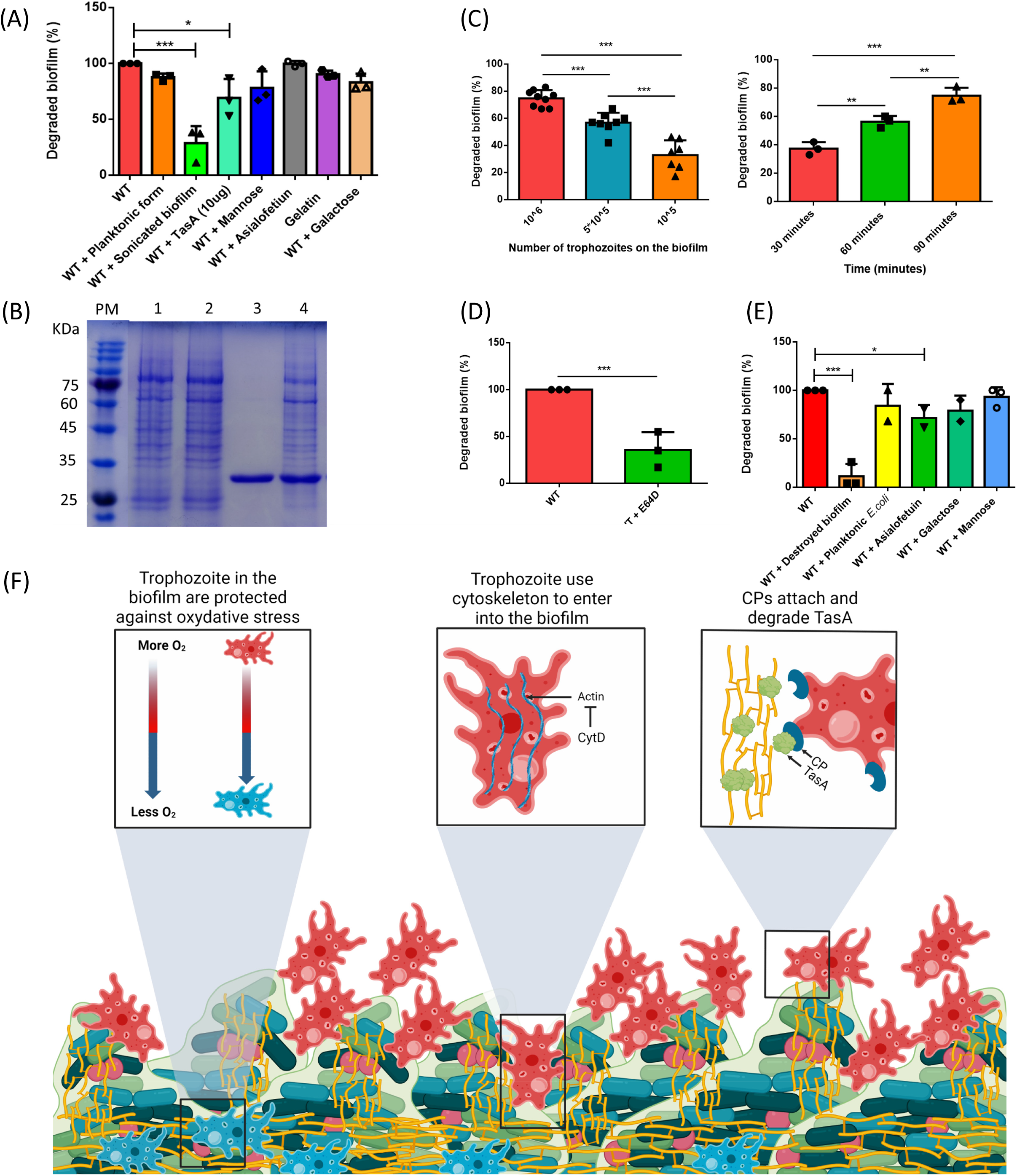
The role of TasA in the predation of *B. subtilis* biofilm cells. (A) Degradation of *B. subtilis* biofilm by *E. histolytica* trophozoites (WT), trophozoites incubated with *B. subtilis* plaktonic form, trophozoites incubated with *B. subtilis* sonicated biofilm, trophozoites incubated with: TasA (10ug), mannose (1%), asialpofetium (0.05%), gelatin (1 %) and galactose (2%). (Student test, ** p-value less than 0.01. *** p-value less than 0.001.). Data represent averages of results from 3 biological replicates. (B) TasA degradation by *E. histolytica* total lysate. (1) Control trophozoites (WT) lysate. (2) TasA (2 ug) + WT lysate (20 ug), (3) TasA and (4) TasA (2 ug) + lysate of E64D treated trophozoites (20 ug). TasA was incubated for 3 hours at 37°C. The degradation of TasA was analyzed by SDS-PAGE following Coomassie staining. (C) *E. coli* biofilm destruction occurs in a time manner. student test, * p-value less than 0.05, ** p-value less than 0.01. *** p-value less than 0.001. Data represent average of results from 3 biological replicates. (D) *E. coli* biofilm destruction occurs in a dose dependent manner. student test, * p-value less than 0.05, ** p-value less than 0.01. *** p-value less than 0.001. Data represent average of results from 3 biological replicates. (E) *E. coli* biofilm degradation after 1 hour of incubation with control trophozoites and with E64D-treated trophozoites. (Student test, ** p-value less than 0.01. *** p-value less than 0.001.) Data represent averages of results from 3 biological replicates. (F) A model representing biofilm-parasite interactions

To test TasA proteinous matrix is also the substrate for CPs from the amoeba, as was suggested by the overall decline in TasA-mCherry fluorescence around the trophozoites (Figure 1A and B), we tested the potential degradation of purified TasA by the extract from trophozoites. We observed that extract from control trophozoites degrade TasA, whereas extracts from E64D-treated trophozoites do not (Fig 4B). Overall, our results suggest that TasA has a dual role in predator prey-interactions with *E. histolytica* serving as the ligand for the binding of the amoeba but also as a target for CPs-mediated degradation. In contrast, it seems that the signal for CP activation is not TasA, as pure TasA failed to induce CPs secretion (Fig. S10).

*E. histolytica* is exposed to a complex bacterial flora in the gut, including biofilm-forming bacteria like *E. coli* EPEC^44^. Our findings that EhCPs can degrade *B. subtilis* biofilm, and the proteinous nature of both *E. coli* and *B. subtilis* biofilms, led us to test whether EhCPs can degrade *E. coli* biofilm. Our data indicate that *E. histolytica* trophozoites are able to degrade *E. coli* biofilm in a time and dose-dependent manner (Figs 4C and D). In contrast, E64D-treated-trophozoites were unable to degrade *E. coli* biofilm (Fig 4E). Based on these results, EhCPs might be involved in the parasite’s general mechanism of bacterial biofilm degradation.

## Discussion

Over the last few decades, it has become obvious that a complex interaction occurs between *E. histolytica* and gut microbiota. This interaction involves dysbiosis of the host’s colonic flora triggered by the ability of the parasite to feed on preferential species of bacteria like *Lactobacillus ruminis* ^45^. Furthermore, some bacteria, such as *E.coli* EPEC O55, regulate the virulence of *E. histolytica* trophozoites and affect their transcriptome ^5^. Many chemical molecules originating from gut microbiota have also an influence on the epigenetic regulation occurring in the parasite and on its resistance to OS and nitrosative stress (for recent reviews on this subject see ^3^ and ^46^). These many studies on the role of the gut microbiota on the biology of *E. histolytica* have focused on the planktonic form of bacteria but none has investigated the biofilm form. The formation of colonic biofilms in healthy individuals is occasional (around 6%), but the presence of these biofilms in individuals with inflamed bowel diseases (IBDs) or with ulcerative colitis (UCs) represent 57% and 34 % of the patients respectively ^47,48^. In healthy individuals, it is assumed that probiotics are the main source of biofilm-forming bacteria. Lactobacillus species like *Lactobacillus rhamnosus* provide a protective shield when forming a biofilm on the tissue of their host ^49^ that can persist there for more than one week after being orally ingested^50^. *B. subtilis* strains, for example, *B. subtilis* PXN21 are a component of a commercial probiotic product, and formulations of probiotics often include spores of *B. subtilis* as their active ingredient ^51^. Traditional foods such as natto contain *B. subtilis* cells and spores and can be used as an alternative to *B. subtilis* probiotics ^52^. While it remains unknown whether these probiotic *B. subtilis* strains are capable of forming biofilms in the GI, the release of biofilm-like microcolonies was shown to enhance the probiotic properties of Firmicutes probiotics^53,54^. Importantly, self-coated biofilms of *B. subtilis* enriched with proteinous extracellular matrix demonstrate extraordinary GI tract tolerance and mucoadhesion, which can substantially improve oral bioavailability and intestinal colonization^54^. In addition to the emerging evidence linking biofilm formation with *B. subtilis* with GI compatibility, numerous pathogens from the Firmicutes phylum form pathogenic biofilms within the gastrointestinal tract. For example, *Enterococcus Faecalis* colonization of the murine GI tract leads to biofilm formation of significant similarity to biofilm formation in vitro^55^. Enteric pathogens were also reported to express biofilm genes in the GI, and matrix proteins. Indeed, patients suffering from IBDs and UCs, dysbiosis of the gut microbiome has been reported with an overgrowth of biofilms formed by *Ruminococcus gnavus* and by EPEC^47^. Therefore, it is highly likely that *E. histolytica* trophozoites will meet bacterial biofilms in the host.

By analogy with the parasite’s interaction with planktonic bacteria and with mammalian cells^56,57^, it is expected that the binding of *E. histolytica* trophozoites to the biofilm represent the first step in the biofilm degradation process. Our data strongly suggest that the Gal/GalNAc lectin^58^ and mannose-receptor^43^ are not involved in the binding process to *B. subtilis* biofilm. Inhibition of *E. histolytica* trophozoites binding to *B. subtilis* biofilm by E64D suggests that CPs are involved in binding. One possible candidate is a 112 kDa adhesin formed by two polypeptides of 49 kDa and 75 kDa that have CP and adhesin activity respectively (for a review see^59^). We also reported that TasA inhibits the binding of *E. histolytica* to *B. subtilis* biofilm, which suggests that TasA is recognized by a TasA binding protein that is present on parasite surface. It is not excluded that CPs and possibly 112 kDa adhesin are involved in the parasite’s binding to the biofilm. TasA serves both as a substrate for trophozoite binding and a target of CPs, suggesting a significant role for matrix proteins in parasite-biofilm interactions. Based on this information, it is also tempting to speculate that TasA triggers EhCP secretion. The lack of induction of EhCPs secretion by pure TasA (Fig. S10), however, refutes this hypothesis.

Following its binding to biofilms, the parasite is expected to degrade it, as it does with planktonic bacteria and mammalian cells ^56,57^. CPs are important virulence factors of *E. histolytica* responsible for the degradation of mucus and extracellular matrix components such as collagen, fibronectin, and immunoglobulins ^60^. Additionally, they facilitate the parasite’s invasion of the colonic mucosa^61^. Of the 20 known *E. histolytica* CPs, three (EhCP1, EhCP2 and EhCP5) account for 90% of the total CP activity ^62^. Our data support a role of *E. histolytica’*s CPs in the degradation of *B. subtilis* and EPEC biofilms. This function of the CPs has also been reported in the unicellular parasite *Giardia lamblia* ^63^which suggests that this is a conserved function among unicellular protozoan. Our work demonstrates that EhCP5 is directly involved in the degradation of *B. subtilis* biofilm. EhCP4 and EhCP6 may also be involved since their expression is upregulated in *E. histolytica* trophozoites exposed to *B. subtilis* biofilm. However, due to the strong sequence homologies between EhCP5, EhCP4 and EhCP6, it is not possible to address the specific role of EhCP4 and EhCP6 by using a sRNA-based RNA interference ^35^. Indeed, precision gene-editing tools like CRISPR-Cas9 are still lacking in *E. histolytica*. This work shows that *E. histolytica’*s CPs are cleaving TasA, a major protein component of the *B. subtilis’* biofilm extracellular matrix ^15,64^. Consequently, TasA degradation by EhCPs can explain the degradation of *B. subtills* biofilms, making it more venerable for trophozoites penetration (Fig. 4F). The degradation of additional biofilm components of *B. subtilis* BslA and TapA by EhCPs is feasible, and may further promote biofilm digestion.

Several studies have shown that membrane traffic and cytoskeletal functions through the modulation of actin dynamics^65^ are critical to the pathogenesis of *E. histolytica*^66,67^. Our study has demonstrated that the degradation of *B. subtilis* biofilms by *E. histolytica* was dependent on the formation of F-filaments, which suggests that mechanical stresses caused by the parasite to deform the biofilm may also contribute to its degradation. The migration of the parasite inside *B. subtilis* biofilm might also be impaired by the inhibition of the parasite’s cytoskeletal functions.

*E. histolytica* degradation of the biofilm is correlated with an upregulation of small GTPase expression. In *E. histolytica* small GTPases are known to regulate actin cytoskeleton, migration, invasion, phagocytosis, and immune response evasion through receptor capping on the surface of the cells ^68^. Among the upregulated small GTPases, 9 encode proteins associated with the AIG1 family. There are 29 known AIG1 family proteins in *E. histolytica* ^69^ but only few have been in depth studied. AIG1 family protein (EHI_176590), which is involved in the formation of protrusions on the plasma membrane and adherence to host cells and AIG1 family protein EHI_127670 which is involved in virulence and controls cysteine peptidase activity of *E. histolytica* ^70^, were not among AIG1 genes upregulated in trophozoites exposed to *B. subtilis* biofilm. The upregulation of many of these AIG1 genes expression suggest their involvement in the degradation of the biofilm by *E. histolytica*.

We found that *E. histolytica* is protected against OS by *B. subtilis* biofilm but not by its planktonic form (Fig. 4F). Oxygen gradient presents inside biofilms create hypoxic zones at their bottom^71^. Upon penetrating the biofilm, the parasite is protected from OS, which promotes its survival. It is also possible that *B. subtilis* biofilms provide to the parasite antioxidant compounds that are not produced by the planktonic form. Activity of tricarboxylic acid (TCA) cycle during early biofilm growth has been reported leading to the accumulation of TCA cycle intermediate including citrate, malate and oxaloacetate ^72^ that are known antioxidants ^73^. One of these compounds, oxaloacetate, protects the parasite against OS ^74^. Interestingly, *B. subtilis* cells seem to also respond to the presence of the parasites. The extract application, and subsequent degradation of the matrix by the parasite significantly induced the cell wall stress response^75^, potentially due to the specific role of TasA as cell-cell adhesion ^33^. The microbial stress from matrix degradation is even more pronounced when measuring a reporter for the general stress response (as judged by the SigB target *ctc*). Canonic environmental stresses activate the stressosome and stress responsive later native sigma factor Sigma B ^76^, however, it is also linked by us to the signaling cascade activated downstream to TasA ^33^. Both cell wall stress response and the general stress response were diminished by E64D. Collectively, these results suggest that *B. subtilis* cells can sense biofilm deterioration and respond accordingly, by inducing their natural defenses associated with stress tolerance. For example, stressosome activation is associated with ROS, low pH and salinity^76^, and cell wall stress response is associated with antibiotic tolerance^75^, all highly relevant to the GI microenvironment.

The multicellular nature of biofilms confers unique phenotypic abilities to the residing bacteria, allowing them to better adhere to, and survive within a host. Therefore, biofilms, and not planktonic cells, are the bacterial entities mostly affecting the human environment. One example is the enormous impact of biofilms on human health. The U.S. Centers for Disease Control and Prevention (CDC) has estimated that bacterial biofilms are responsible for 60% of chronic infections, including burn wounds, chronic ulcers of limbs associated with diabetes, periodontitis, osteomyelitis, chronic wounds, and cystic fibrosis lungs^24,77^. Biofilm cells are inherently more resistant to the host immune system and to antibiotics^78^. The tolerance of infection-forming biofilms to antibiotics is reported to 10- to 1,000-fold less susceptible to antibiotics than their planktonic counterparts^79^. Thereof, a new reservoir of anti-biofilm approaches is vital. So far, anti-biofilm agents based on natural products included primarily phytochemicals, such as phenolics, essential oils, terpenoids, lectins, alkaloids, polypeptides, and polyacetylenes targeting microbial communication and/or adhesion, biosurfactants, reducing surface-bacteria interactions, and quorum sensing inhibitors of natural source, directly targeting microbial signal pheromones^80^. Enzymes were also suggested as anti-biofilm strategies with clinical relevance, and were considered to be produced in natural scenarios during competition between bacterial biofilms^81^. One unexplored resource for uncovering novel anti-biofilm agents with a broad spectrum are protozoan parasites. This neglect is not trivial as single-cells eukaryotes and bacteria exert well-established predator-prey interactions ^82^, which should be extendable to bacterial biofilms in various ecological niches, most famously, zooplankton grazers-phytoplankton mats interactions^83^. Here we provided a proof of concept that indeed *E. histolytica* parasites were evolved to specifically interact with, and destroy biofilms of two phylogenetically unrelated bacteria, *B. subtilis* and *E. coli*, both robust biofilm formers. The specific activation of matrix degrading CPs in response to the microbial biofilm is strongly suggesting that amoeba are adapted to biofilm preys, and may serve as a new unexplored reservoir of novel therapeutic approaches to treat biofilms. Furthermore, our findings here that *E. histolytica* trophozoites can use the biofilm as a shield reducing the oxidative stress of the parasite highlight biofilm-amoeba interactions as one unexpected significant regulator of the parasite’s stress tolerance and pathogenicity.

## Methods

### Culture of *E. histolytica*

*E. histolytica* trophozoites, the HM-1:IMSS strain (from Prof. Samudrala Gourinath, Jawaharlal Nehru University, New Delhi, India), were grown at 37 °C in 13 × 100 mm in screw-capped Pyrex glass tubes in serum free Diamond’s TYI S-33 medium (Johnson and Johnson, Hyclone, USA) to exponential phase. Trophozoites were harvested from their growth support by incubating the tubes by tapping the glass tubes followed by centrifugation (Eppendorf centrifuge 5810R, rotor A-4-62) according to a previously reported protocol ^84^.

### Biofilm formation

#### B.subtilis

All experiments were performed with *B. subtilis* GFP-expressing strain NCIB3610, amyE::P hyperspank-gfp. A single colony were isolated from lysogeny broth (LB) plates were grown to mid-logarithmic phase in a 3-ml LB culture (4 hours at 37°C with shaking, 200 rpm, New Brunswick scientific, Innova 4300). For biofilms preparation, cells from mid-logarithmic phase were diluted 1:10 into serum free Diamond’s TYI S-33 medium and grown in 24-well plates cover with 1mL of serum free Diamond’s TYI S-33 agar as shown in Xiaoling Wang and al, 2015 ^85^ at 30°C, without shaking overnight.

#### E. coli

All experiments were performed with *E.coli* GFP-expressing strain MG1655 (. A single colony were isolated from lysogeny broth (LB) plates were grown to mid-logarithmic phase in a 3-ml LB broth culture (4 hours at 37°C with shaking, 200 rpm). For biofilms preparation, cells from mid-logarithmic phase were diluted 1:10 into serum free Diamond’s TYI S-33 medium and grown in 24-well plates at 37°C, without shaking overnight.

### Degradation of *B. subtilis/E.coli* biofilm by *E. histolytica* trophozoites

Trophozoites (1×10^6^) were incubated on the biofilm (37 °C for *B.subtilis* 3 hours for *B.subtilis* and 1 hour for *E.coli*, without shaking). After incubation, trophozoites were removed and the biofilm was washed one time with serum free Diamond’s TYI S-33 medium. The quantification of biofilm degradation was done by comparing the GFP expression level of each well to the control (biofilm incubated without trophozoites). The pictures of each biofilm were taken with the Olympus MVX10 (Olympus DP73 camera, GFP laser and 2.5 zoom) and the software cellsens dimension. The picture’s GFP signal was analyzed with FIJI ImageJ.

### RNA extraction

Trophozoites (10^6^ cells/ml) were incubated (1 hours at 37°C) with *B. subtilis* planktonic form, *B. subtilis* biofilm form or with serum free Diamond’s TYI S-33 medium without bacteria as control, in 24 well plate covered with serum free Diamond’s TYI S-33 agar. After incubation, trophozoites were harvested by tapping the glass tube followed by a centrifugation (1900rpm, 3 minutes) *E. histolytica* RNA was prepared using the Monarch Total RNA Miniprep Kit (NEW ENGLAND BioLabs, Ornat, Nes Ziona, Israel). According to the manufacturer’s instructions, the absence of mechanical disruption during the cell lysis step favors the extraction of *E. histolytica* RNA over *B.subtilis* biofilm RNA using the detergent-based lysis buffer.

### Flow cytometry

Cells were harvested from established pellicle biofilms grown in MSgg in the presence or absence of indicated extracts. Biofilms were mildly sonicated. For flow cytometry analysis, cells were suspended in PBS and measured on a BD LSR II flow cytometer (BD Biosciences). The GFP fluorescence was measured using laser excitation of 488 nm and 561 nm (respectively), coupled with 505 LP and 525/50 sequential filters. The photomultiplier voltage was set to 484 V. 100,000 cells were counted, and every sample was analyzed with Diva 8 software (BD Biosciences).

### Scanning Electron Microscopy

Biofilms grown as described above with the trophozoite’s over a mesh. The mesh was gently removed with the intact biofilms and samples fixed for 2-4 hours at 4°C with 2% glutaraldehyde, 3% paraformaldehyde, 0.1 M sodium cacodylate (pH 7.4) and 5 mM CaCl2. After two 15 min washes with double distilled water, samples were dehydrated through series of ethanol washes. Subsequently, samples were dried overnight at room temperature. The samples were sputter-coated with gold–palladium shortly before examination with a scanning electron microscope (SEM) XL30 with field emission gun.

### Library construction and sequencing

The library construction and deep sequencing were done as described in ^86^. Briefly, night RNAseq libraries were produced according to the manufacturer’s protocol (NEBNext UltraII Directional RNA Library Prep Kit, Illumina, NEB, MA, USA) using aroung 1 000 ng of total RNA. The RNAseq data was generated on an Illumina NextSeq500, 75 single-end read, high output mode (Illumina).

### Virulence factors and *B. subtilis* biofilm degradation

Trophozoites (1×10^6^) were incubated with the actin polymerization inhibitor cytochalasin D (CytD) (5 μM), or the cysteine proteinase inhibitor E64D (10 μM) for 24h at 37°C. The viability of the trophozoites was measured by using eosin exclusion assay^87^. For each condition, 10^6^ trophozoites were incubated with the biofilm at 37 °C for 3 hours. We used trophozoites fixed with paraformaldehyde, (4%) as a negative control. After incubation, trophozoites were removed by washing once the wells with serum free Diamond’s TYI S-33 medium. Biofilm degradation for each condition was measured as described above.

### Virulence factors and *E. coli* biofilm degradation

Trophozoites (1×10^6^) were incubated with the cysteine proteinase inhibitor E64D (10 μM) for 24h at 37°C. The viability of the trophozoites was measured by using eosin exclusion assay^87^. For each condition, 10^6^ trophozoites were incubated with the biofilm at 37 °C for 1 hours. After incubation, trophozoites were removed by washing once the wells with serum free Diamond’s TYI S-33 medium. Biofilm degradation for each condition was measured as described above.

### Competition with *B. subtilis* biofilm

Trophozoites (1×10^6^) were incubated with the biofilm (for 3 hours at 37°C) with galactose (2%, Acro organic), mannose (1%, Acro organic), asialofetium (0.05%, Sigma), gelatin (1%, Sigma), TasA (10μg), planktonic form (1×10^9^) and sonicated biofilm (one biofilm was sonicated for 3 cycles of 15 s on/off at medium intensity on abioruptor UCD 200) of *B. subtilis* as competitor. After incubation, trophozoites were removed and the biofilm was washing one time the well with serum free Diamond’s TYI S-33 medium. Biofilm degradation was determined by measuring the GFP signal level that remains following the action of *E.histolytica* trophozoites and the data were normalized to the GFP signal of biofilm incubated without trophozoites. The level of GFP signal in each biofilm was measured with an Olympus MVX10 (Olympus DP73 camera, GFP laser and 2.5 zoom) and the picture were treated with the software cellsens dimension and with FIJI imajeJ.

### Immunofluorescence inverted confocal microscopy

Trophozoites (1×10^6^) were incubated with *B.subtilis* biofilm for 30 minutes or 180 minutes at 37°C. Trophozoites that did not attached were wash away with serum free Diamond’s TYI S-33 medium. The biofilm was fixed with paraformaldehyde (4%, 30 minutes at RT) and stain with DAPI (20 μg/ml, 1 hour at 4°C in the dark). The biofilm was transferred to microscope slides and were then examined under an inverted confocal immunofluorescence microscope (Zeiss LSM700 meta laser scanning confocal imaging system, zoom 200). For analysis, the number of trophozoites on biofilm were counted and biofilm destruction was quantified for each conditions using Imaris software.

### Quantitative PCR

Quantitative amplification was performed as mention in^88^, on WT and siEhCP5 trophozoites gDNA. The list of primer used in this study is described in table 1.

### Silencing of EhCP5 (EH_168240) gene, transfection

For the construction of the siEhCP5 silencing vector that was used to silence EhCP5 expression, EhCP5 (EH_168240) was amplified from *E. histolytica’*s genomic DNA using the primers 5’EhCP5 and 3’EhCP5 (table 1). The resulting PCR product was cloned into the pGEM-T Easy vector system (Promega, WI, USA) and then digested with the restriction enzymes, BglII and XhoI. The digested DNA insert was subcloned into the *E. histolytica* pEhEx-04-trigger silencing vector, containing a 142-bp trigger region (EHI_048660), from Tomoyoshi Nozaki, University of Tokyo, Japan, to generate siEhCP5 vector.

### Cysteine proteinase (CPs) activity assay

CPs activity was measured in total lysates of trophozoites (1×10^6^) lysed in 1 ml of Nonidet P-40 (1% in phosphate-buffered saline, PBS). CPs activity was measured according to ^89^. One unit of activity corresponds to the number of micromoles of Z-Arg-Arg-pNA (BACHEM) digested per min per mg of protein.

### Detection of oxidized proteins

The level of oxidized protein was measured with the Oxyblot kit (Protein Carbonyl Assay Kit, Abcam) according to the manufacturer protocol. Trophozoites (1×10^6^) were incubated for 1 hours at 37°C with *B. subtilis* planktonic form, *B. subtilis* biofilm form or with serum free Diamond’s TYI S-33 medium without bacteria (as control) in 24 well plates cover with serum free Diamond’s TYI S-33 agar. Next, the cells were exposed to H_2_O_2_ (2.5 mM, for 30 minutes at 37°C). Trophozoites were then lysed with Nonidet P-40 (1% in phosphate-buffered saline, PBS) for 15 minutes on ice. Equal protein concentrations (20 μg) were proceed with the oxyblot, protein oxidation detection kit ^90^.

### TasA purification

Protein was expressed and purified as previously described^91^. TasA was purified using the pDFR6 (pET22b-*tasA*) and *E. coli* BL21(DE3) cells were freshly transformed with the plasmid. Colonies were selected from the plates and resuspended in 10 mL of LB with 100 μg/mL of ampicillin and incubated overnight at 37 °C with shaking. This pre-inoculum was then used to inoculate 500 mL of LB + ampicillin, and the culture was incubated at 37 °C until an OD600 of 0.7–0.8 was reached. Next, the culture was induced with 1-mM isopropyl β-D-1-thiogalactopyranoside (IPTG) and incubated O/N at 30 °C with shaking to induce the formation of inclusion bodies. After that, cells were harvested by centrifugation (5000 × g, 15 min, 4 °C) resuspended in buffer A (Tris 50 mM, 150 mM NaCl, pH8), and then centrifuged again. These pellets were stored frozen at −80 °C until used. After thawing, cells were resuspended in buffer A, and broke down by sonication on ice using a Branson 450 digital sonifier (3 × 45 s, 60% amplitude). After sonication, the lysates were centrifuged (15,000 × g, 60 min, 4 °C) and the supernatant was discarded, as proteins were mainly expressed in inclusion bodies. The proteinaceous pellet was resuspended in buffer A supplemented with 2 % Triton X-100, incubated at 37 °C with shaking for 20 min., to further eliminate any remaining cell debris, and centrifuged (15,000 × g, 10 min, 4 °C). The pellet was then extensively washed with buffer A (37 °C, 2 h), centrifuged (15,000 × g for 10 min, 4 °C), resuspended in denaturing buffer (Tris 50 mM NaCl 500 mM, 6 M GuHCl), and incubated at 60 °C overnight to completely solubilize the inclusion bodies. Lysates were clarified via sonication on ice (3 × 45 s, 60% amplitude) and centrifugation (15,000 × g, 1 h, 16 °C) and were then passed through a 0.45-μm filter prior to affinity chromatography. Proteins were purified using an AKTA Start FPLC system (GE Healthcare). The lysates were loaded into a HisTrap HP 5 mL column (GE Healthcare) previously equilibrated with binding buffer (50 mM Tris, 0.5 M NaCl, 20 mM imidazole, 8 M urea, pH 8). Protein was eluted from the column with elution buffer (50 mM Tris, 0.5 M NaCl, 500 mM imidazole, 8 M urea, pH 8). After the affinity chromatography step, buffer was exchanged to 1% acetic acid pH 3, 0.02% sodium azide by using a HiPrep 26/10 desalting column (GE Healthcare). This ensured that the proteins were maintained in their monomeric form. The purified proteins were stored under these conditions at 4 °C (maximum 1 month) until further use.

### TasA degradation by trophozoites lysate

Trophozoites were lysed with Nonidet P-40 (1% in phosphate-buffered saline, PBS) for 15 minutes on ice. Then, TasA (2μg) was incubated with different concentration of trophozoites lysate (5 μg, 10 μg, 15 μg and 20 μg) for 1 hour and 3 hours at 37°C in a final volume of 20 μl with DTT (1M). Different conditions were analyzed by 12% SDS page followed by Coomassie staining. As a control TasA and trophozoites lysate were incubated alone. The quantification was done with FIJI ImageJ.

## Notes

### Competing Interest Statement

The authors have declared no competing interest.

