## Supporting Figures and Tables for "Digestive exophagy of Bacterial Biofilms by an Amoeba Predator is Mediated by Specific Biofilm Recognition"

### Supporting Information

Supporting Figures (S1-S10)

Supporting tables (1-3)

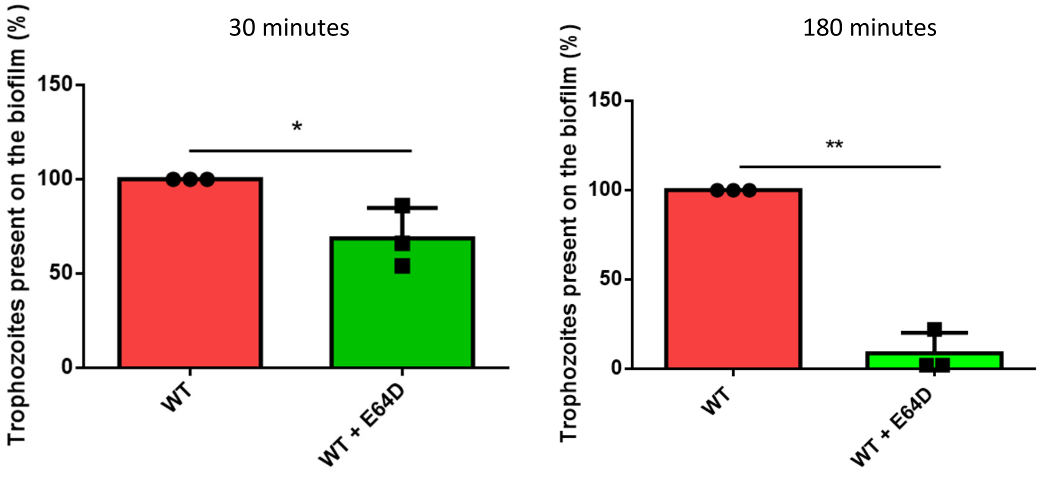


**Figure S1.** Quantification with Imaris of the number of trophozoites found on the surface or inside the biofilm after 30 minutes and 180 minutes of incubation at 37°C. Trophozoites non-treated with E64D are taken as control. student test, * p-value less than 0.05. Data represent average of results from 3 biological replicates.

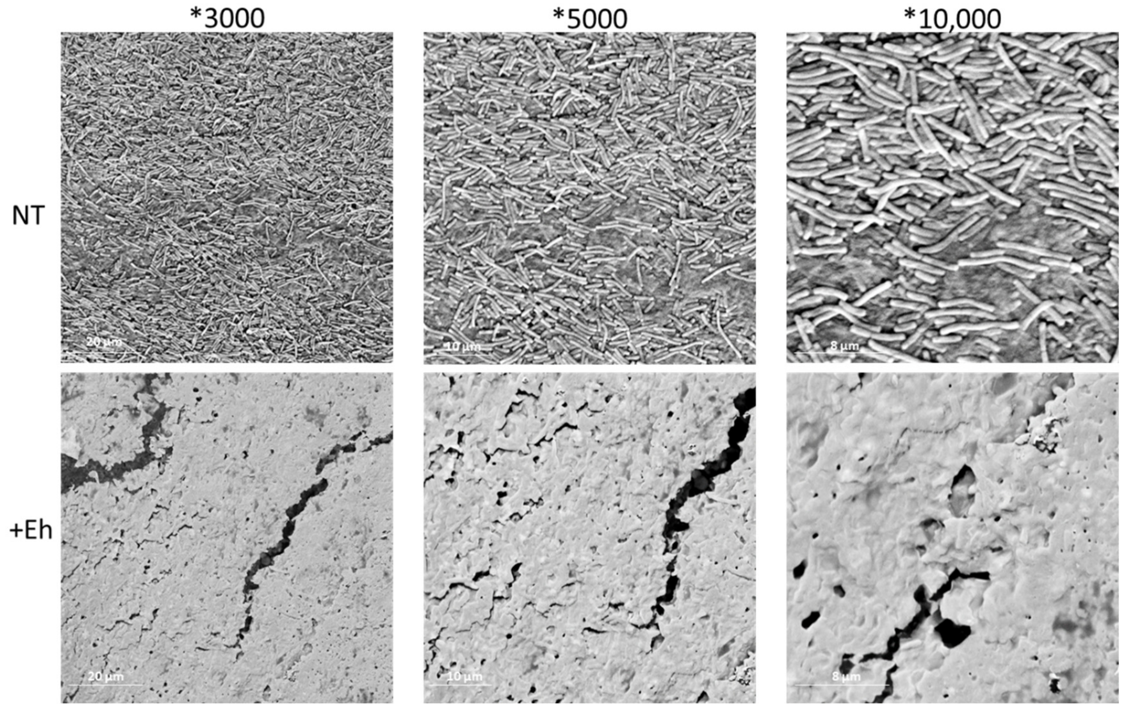


**Figure S2.** Scanning Electron Microscopy of biofilm surfaces of treated and untreated biofilms for 3 hours grown as in Figure 1A. Shown are different magnifications of representative fields

Planktonic versus control

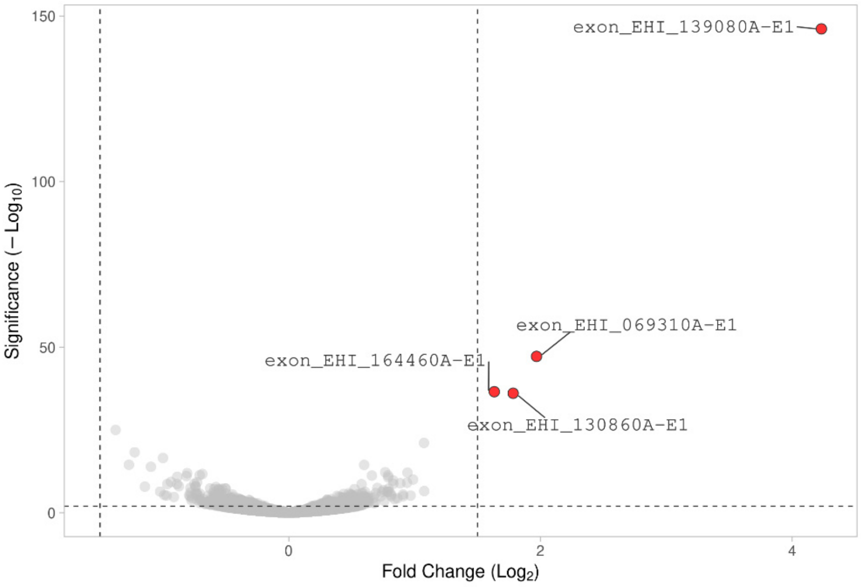


Biofilm vs. control

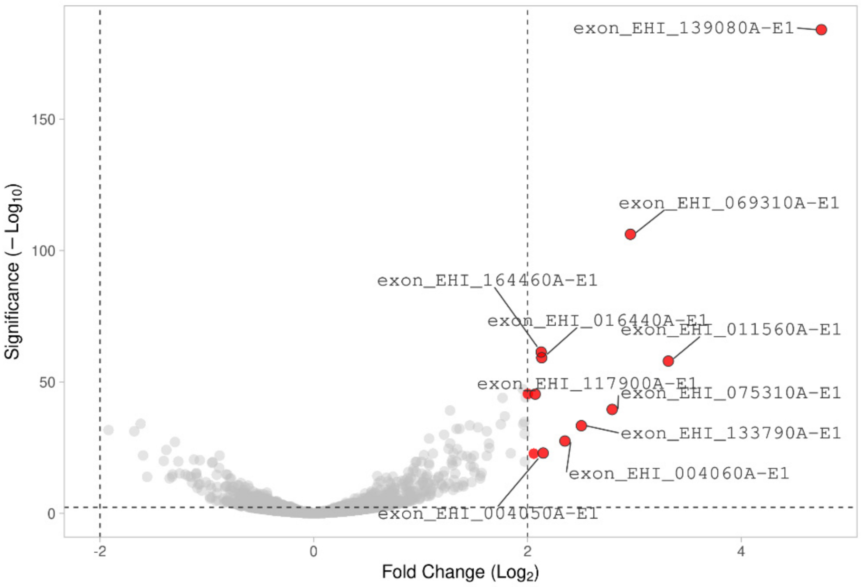


**Figure S3.** Volcano plots. The red dots represent differentially expressed genes between the indicated treatments. The other genes are represented by the light gray dots.

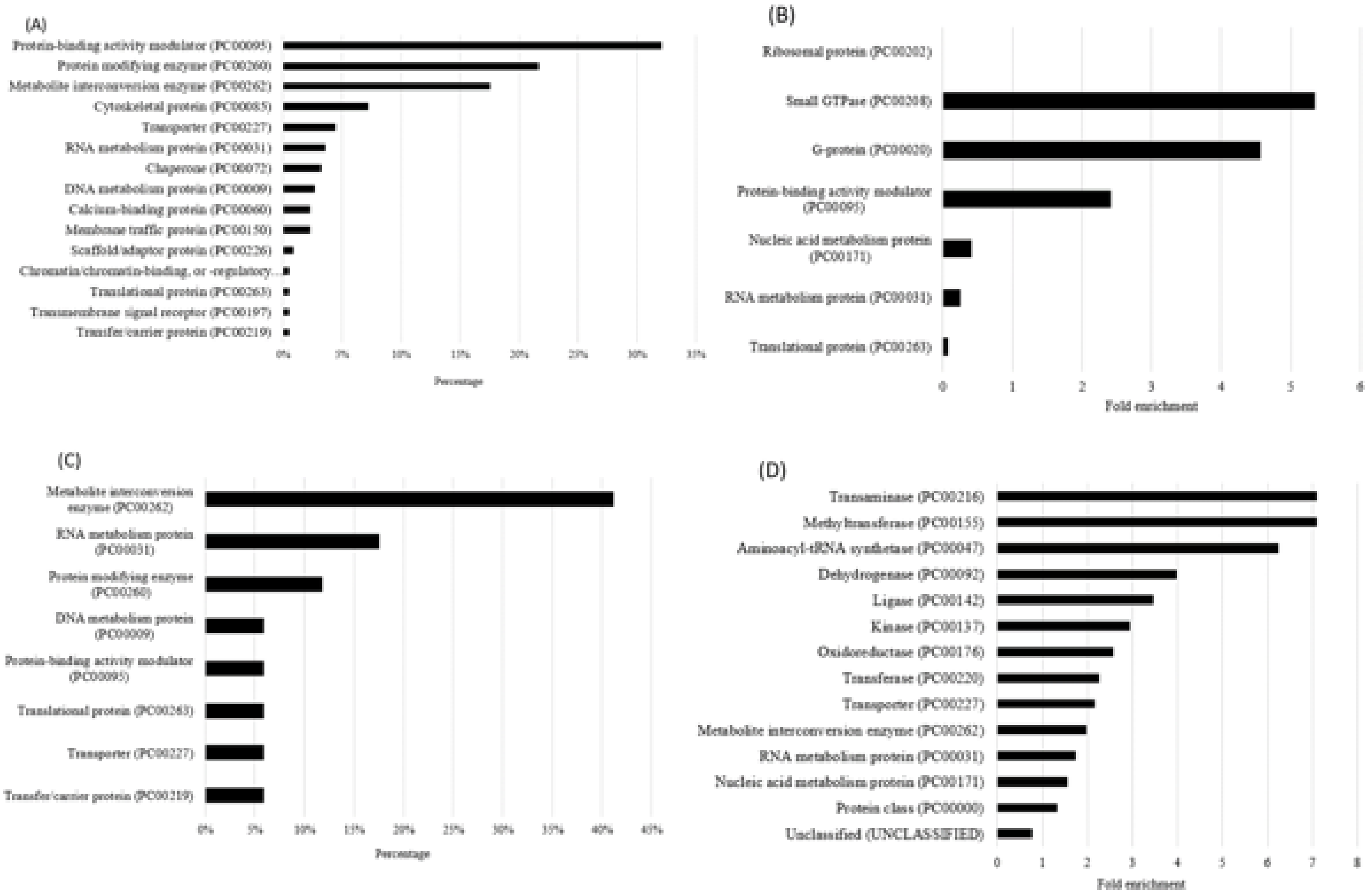


**Figure S4.** Transcriptomic results of trophozoites incubated with Planktonic form of B. subtilis VS control trophozoites. (A) PANTHER sequence classification of genes upregulated in AFAT. (B) PANTHER statistical overrepresentation test of upregulated genes in AFAT; (C) PANTHER sequence classification of genes downregulated in AFAT. (D) PANTHER statistical overrepresentation test of downregulated genes in AFAT.

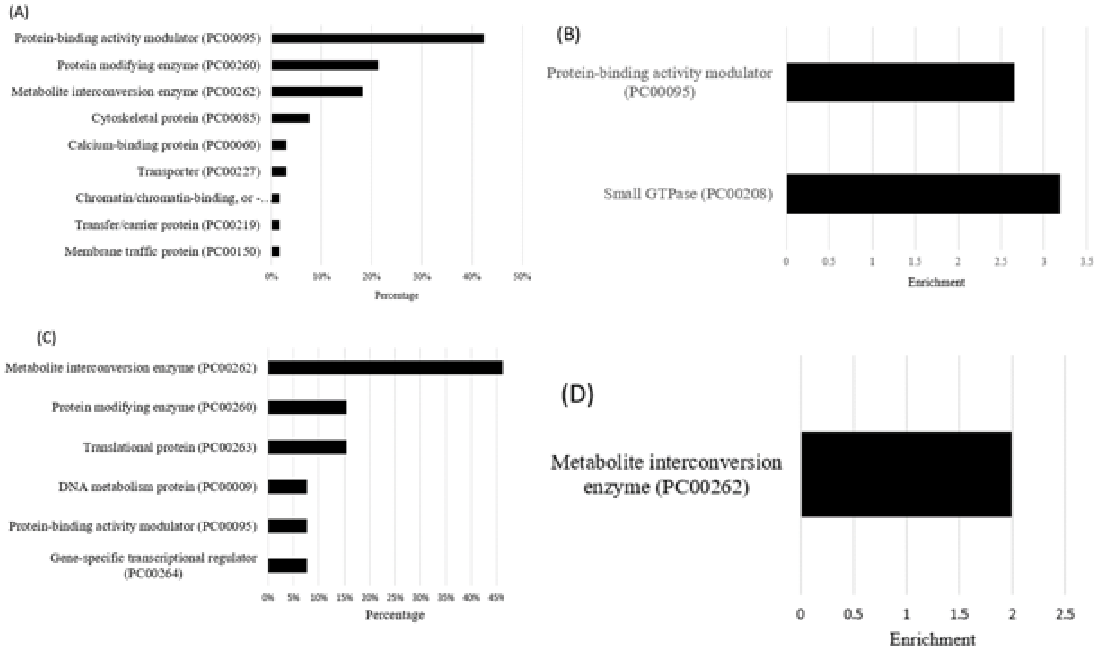


**Figure S5.** Transcriptomic results of trophozoites incubated with Planktonic form of B. subtilis VS control trophozoites. (A) PANTHER sequence classification of genes upregulated in AFAT. (B) PANTHER statistical overrepresentation test of upregulated genes in AFAT; (C) PANTHER sequence classification of genes downregulated in AFAT. (D) PANTHER statistical overrepresentation test of downregulated genes in AFAT.

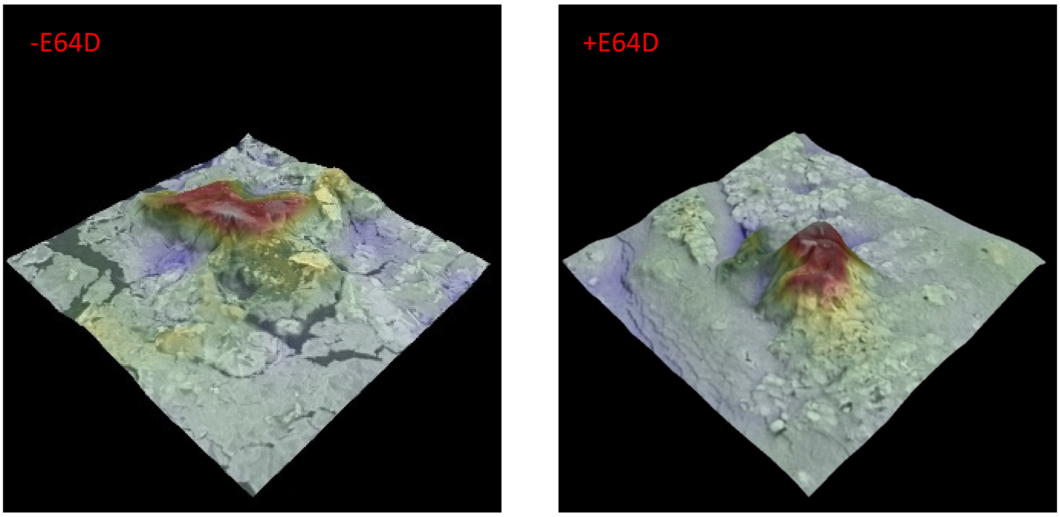


**Figure S6.** An heat map of thickness of biofilms (gradients are from low-blue, to high- red) analyzed by scanning electron microscopy (Figs 1A and B) demonstrates the parasite footprint as well as areas of clearance in biofilms treated with trophozoites in the absence of E64D but not its presence.

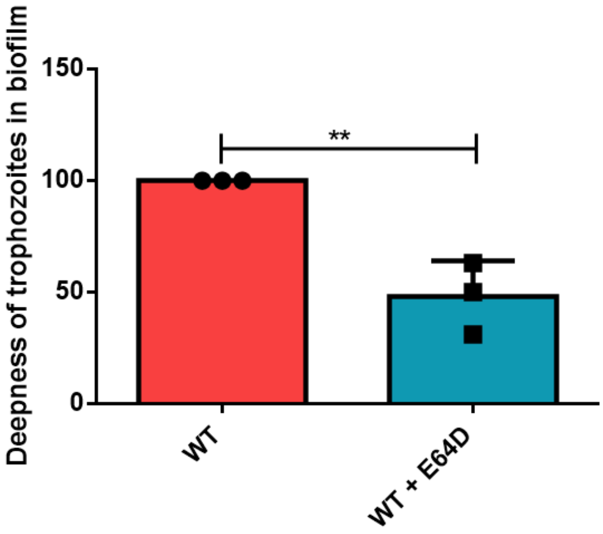


**Figure S7.** Quantification with Imaris of the deepness on trophozoites treated or not with E64D in the biofilm after 180 minutes. Trophozoites entered inside in the biofilm at a depth of 10 to 40 µm. Student test, * p-value less than 0.05. Data represent average of results from three biological replicates.

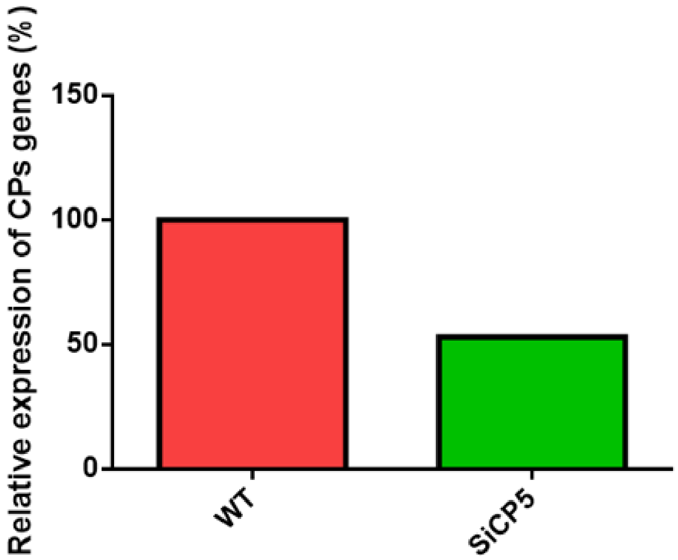


**Figure S8:** qRT-PCR analysis of EhCP5 expression in control trophozoites and in siEhCP5 trophozoites EhCP5 (Eh_5168240) expression levels in E. histolytica trophozoites. The relative fold change of EhCP5 expression in control (WT) and siCP5 trophozoites was calculated using the 2 ^(−ΔΔCt) method 91. Data are from one biological replicate, each with three technical replicates.

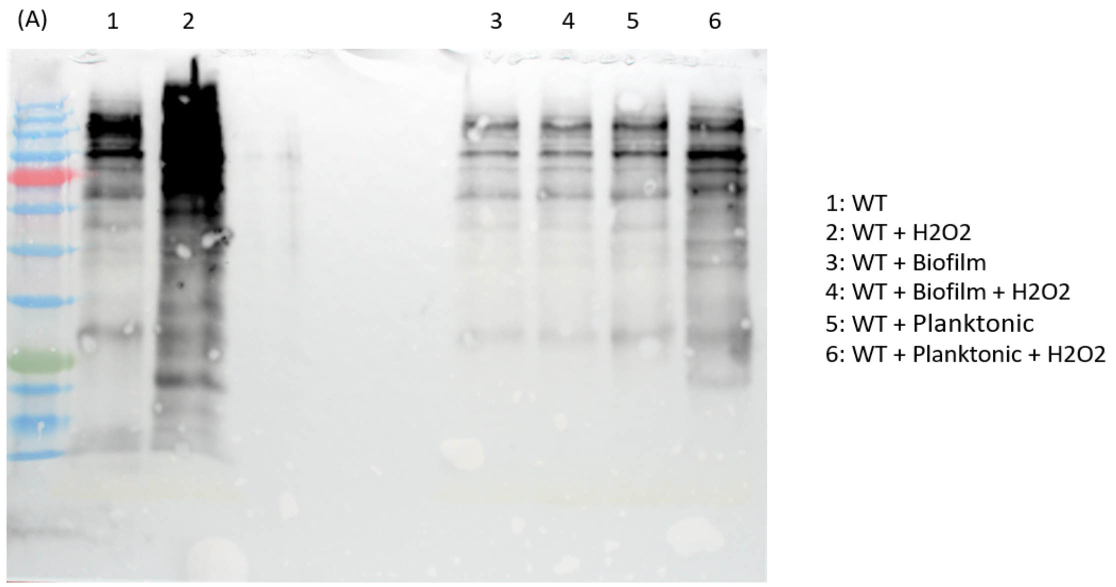


**Figure S9:** Western-blot analysis of oxidized proteins in trophozoites incubated with planktonic or biofilm *B. subtilis* and exposed to H2O2 (2.5 mM, 30 min).

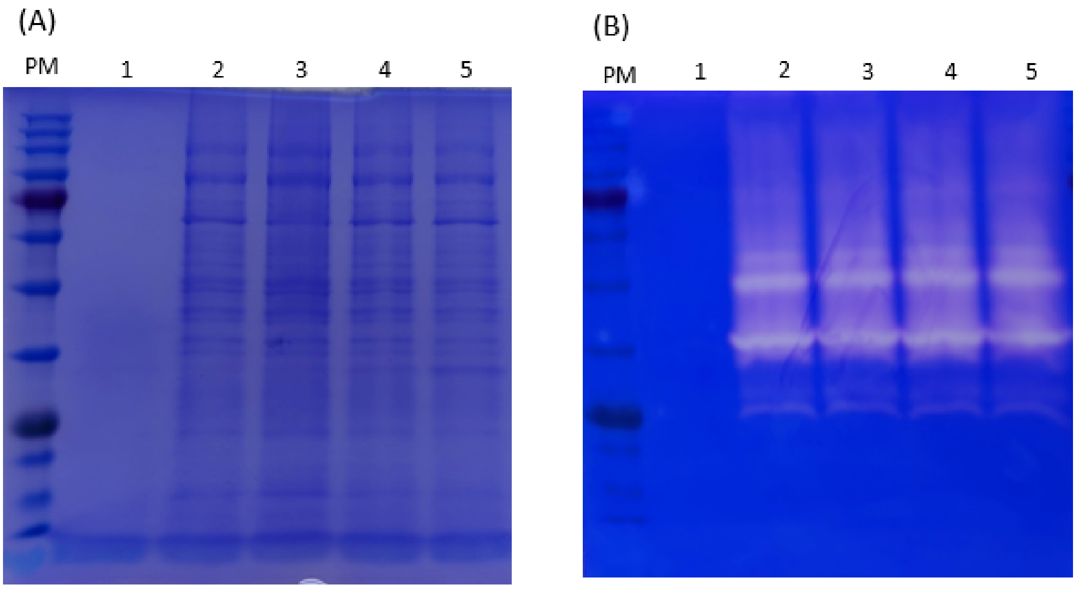


**Figure S10.** TasA does not boost the secretion of EhCPs. TasA and trophozoites secretion product (SP) were incubated 3 hours at 37°C in 500 µL of YI_secretion medium (as described in 92). Protein were separated on (A)12% SDS-Page or on (B) a gelatin gel (1% gelatin) followed by staining with coomassie. Legend: (PM) protein marker, (1) control TasA, (2) control SP, (3) SP + 2 ug TasA, (4) SP + 5 ug TasA and (5) SP + 10 ug Ta.sA.

**Supporting Tables**

**Supporting Table 1: Primers used for Q-PCR**

| Gene Name | Gene symbol | Oligonucleotide sequence 5’ | Oligonucleotide sequence 3’ |
|---|---|---|---|
| EhCP4 | EHI_168240 | CCAGAATCTGTTGATTG GAGA | GCAACCAACAATCTTCCTTC |
| EhCP5 | EHI_050570 | CAGAAGGACCAGTTGCT GTT | ATATCCTACAGCGGCAACA C |
| EhCP6 | EHI_151440 | TTGCTATTGATGCAGGT CAA | AGATCCATATCCAACAGCAC A |
| Actin | EHI_142730 | TTAACTGAAAGAGGATAT GCT | T TCACTGCTTGATGCAGCTTT TTG |

**Supporting Table 2: Transcriptomics of WT trophozoites vs. WT trophozoites incubated with planktonic *B. subtilis* vs. WT trophozoites incubated with *B. subtilis* biofilms.** Results are attached as an independent Excel file titled "Supporting Table 2")

**Supporting Table 3: List of AIG1 genes upregulated when *E. histolytica* trophozoites are incubated with *B. subtilis* biofilm.**

| AIG1 gene | FC |
|---|---|
| EHI_136950 | 2.1 |
| EHI_126560 | 1.4 |
| EHI_176590 | 1.5 |
| EHI_136940 | 2.4 |
| EHI_102600 | 1.6 |
| EHI_119040 | 1.6 |
| EHI_089670 | 1.6 |
| EHI_195260 | 2 |
| EHI_144270 | 1.7 |
| EHI_176580 | 1.8 |
